# Enriching hippocampal memory function in older adults through real-world exploration

**DOI:** 10.1101/586974

**Authors:** Branden S. Kolarik, Shauna M. Stark, Samantha M. Rutledge, Craig E.L. Stark

## Abstract

Age-related structural and functional changes in the hippocampus can have a severe impact on hippocampally-dependent memory performance. Here we test the hypothesis that a real-world spatial exploration intervention will improve hippocampally-dependent memory performance in healthy older adults. We found that following our intervention, participants’ lure discrimination index (LDI) was significantly higher than it was at baseline, while traditional recognition scores remained relatively unchanged. These results point to the viability of a spatial exploration intervention for improving hippocampally-dependent memory in older adults.

## 1.1 Introduction

Environmental enrichment provides animals with an environment, which allows for physical exercise and exploration. Exposure to an enriched environment can promote neurogenesis, synaptogenesis and improve performance on hippocampal-dependent memory tasks (Kempermann, Kuhn, & Gage, 1997; van Praag, Kempermann, & Gage, 1999; Clemenson et al., 2015). One important component of environmental enrichment appears to be the amount spatial exploration of the environment. Animals that spend more time exploring their environment show increased neurogenesis in the dentate gyrus of the hippocampus (Freund et al., 2013). In humans, extensive spatial exploration training has been linked to increased hippocampal volume in both real (Maguire et al., 2000) and virtual environments (Lövdén et al., 2012). Normal healthy aging can also bring about structural and functional changes in the hippocampus (Raz et al., 2005; O’Shea, Cohen, Porges, Nissim, & Woods, 2016) and can have a profound impact on hippocampal-dependent memory performance (Yassa, Mattfeld, Stark, & Stark, 2011). In animals, however, due the plastic nature of the hippocampus, these changes can be partially alleviated through environmental enrichment (Bennett, Mcrae, Levy, & Frick, 2006; Kempermann, Kuhn, & Gage, 1998). Given our understanding of age-related structural and behavioral changes, as well as the impact of environmental enrichment on the hippocampus, how can we promote memory improvement in humans with environmental enrichment?

One possible method for promoting memory improvement in humans is the use of video games as a form of environmental enrichment. Video games, particularly 3D video games that allow for spatial exploration of a virtual environment, are a way for humans to interact with a novel, enriched environment and learn the spatial layout of landmarks and goal locations.

Previous work from our lab, using video games as a behavioral intervention, showed that playing games for 30 minutes per day for two weeks improves hippocampal-based mnemonic pattern separation (Clemenson, Henningfield, & Stark, 2019; Clemenson & Stark, 2015). Importantly, this effect was present specifically for those who played a rich 3D rather than a simple 2D video game (Clemenson & Stark, 2015) and correlated with their amount of spatial exploration (Clemenson et al., 2019). These results suggest that the spatial exploration of a virtual environment can promote changes in hippocampally-dependent memory tasks, similar to what has been seen previously in rodents.

In the current study, we set out to investigate if a real-world spatial exploration task, similar to that used in the video game intervention, can also have an impact on hippocampal-based memory function. We developed a scavenger hunt task that participants performed over the course of a four-week behavioral intervention. We assessed hippocampal memory function with the Mnemonic Similarity Task (MST) (Kirwan & Stark, 2007; Stark, Yassa, Lacy, & Stark, 2013) before and after our intervention as well as at a four-week follow-up test. We focused on the MST’s lure discrimination index (LDI), which tests object recognition memory using highly similar lure items in an effort to tax the hippocampal function known as pattern separation (Rolls, 2016; Yassa & Stark, 2011). The MST LDI has been shown to be highly sensitive to hippocampal function (Kirwan & Stark, 2007; Stark et al., 2013), to age-related memory decline (Holden, Toner, Pirogovsky, Kirwan, & Gilbert, 2013; Stark & Stark, 2017; Stark, Stevenson, Wu, Rutledge, & Stark, 2015; Stark et al., 2013; Toner, Pirogovsky, Kirwan, & Gilbert, 2009) and to hippocampal connectivity (Bennett, Huffman, & Stark, 2015; Bennett & Stark, 2015; Yassa et al., 2011; Yassa, Muftuler, & Stark, 2010) and function. Since environmental enrichment and spatial exploration have been shown to specifically affect the hippocampus in rodent models, we hypothesized that our real-world exploration intervention should have the most effect on hippocampal-dependent memory (MST LDI) and little effect on traditional recognition memory.

Here, we show that mnemonic discrimination (MST LDI) significantly improved following our intervention and that scores remained higher than pre-test baseline performance, even four weeks after stopping the intervention, while recognition scores were relatively unaffected. This observed change in memory ability was not related to pre-test baseline scores, the rate of learning in our behavioral intervention, or the amount of physical activity completed during the intervention, indicating that the changes can likely be attributed to the spatial exploration and learning that occurred during the task.

## 2 Method

### 2.1 Participants

Twenty-six adults between the ages of 65-79 were recruited from the community around UC Irvine and screened for cognitive impairments prior to inclusion in the study. One participant dropped out before completing the study, so our final sample included 25 adults (mean age = 69.8, 20 Female). All participants were compensated for their participation in the study and provided informed consent in accordance with the University of California, Irvine Internal Review Board.

### 2.2 Experimental Procedure

#### 2.2.1 Neuropsychological Tests

Participants were given a battery of standardized neuropsychological tests to ensure their cognitive abilities were within normal range for their age: Mini Mental State Exam (Crum, Anthony, Bassett, & Folstein, 1993), Geriatric Depression Scale (Yesavage et al., 1982), Rey Auditory Verbal Learning Test (Rey, 1941), Rey-Osterrieth Complex Figure (Osterrieth, 1944), Trails A and B (Tombaugh, 2004), Letter Number Sequencing (Wechsler, 1997), Digit Span (Wechsler, 1997), and Stroop Task (Stroop, 1935) (Table 1).

**Table 1.** Neuropsychological test score means (sd). MMSE = Mini Mental State Exam, GDS = Geriatric Depression Scale, RAVLT = Rey Auditory Verbal Learning Test, LN Seq = Letter Number Sequencing

|  |  |
| --- | --- |
| Age | 69.9 (4.1) |
| Education | 14.8 (2.6) |
| MMSE | 28.7 (1.0) |
| GDS | 1 (1.71) |
| RAVLT Total | 49.0 (9.3) |
| RAVLT Immediate | 10.6 (3.1) |
| RAVLT Delay | 10.8 (3.4) |
| Rey-O Figure | 35.0 (1.4) |
| Rey-O Delayed | 18.6 (5.1) |
| Trail A | 31.0 (9.8) |
| Trail B | 76.1 (25.2) |
| Digit Span Forward | 10.6 (2.4) |
| Digit Span Backward | 7.6 (1.8) |
| Digit Span Total | 18.2 (3.8) |
| LN Sequence | 11.4 (2.4) |
| Stroop W T Score | 44.5 (11.4) |
| Stroop C T Score | 46.5 (10.6) |
| Stroop CW T Score | 48.6 (8.3) |
| Stroop Inter T Scores | 44.6 (6.7) |

#### 2.2.2 Mnemonic Similarity Task

As part of the behavioral measures, participants completed the Mnemonic Similarity Task (Stark et al., 2013) (Figure 1A) at three separate time points (pre-intervention, post-intervention, and follow-up). During the incidental encoding phase, 128 everyday items were presented on the screen, one at a time, for a total of 2.5 seconds each (0.5 s ISI). Participants made judgments of whether each item was an “indoor” or “outdoor” item by tapping the appropriate response button on a touchscreen laptop. Immediately following encoding, participants performed a recognition memory task for 192 items (64 repeated items, 64 lure items, and 64 foil items) presented for 2.5 seconds each (0.5 s ISI). Of the 192 items, 64 were repeats from the encoding phase (targets), 64 were similar, but not identical, to an image seen during encoding (lures), and 64 were new (foils). Trial types were presented randomly. Participants again made a response on a touchscreen laptop to judge whether an item was “Old”, “Similar”, or “New”. A lure discrimination index (LDI) was calculated as the difference between the rate of “Similar” responses given to lure items minus the rate of “Similar” responses given to foils. A traditional recognition memory measure was calculated as the difference between the rate of “Old” responses to repeats minus “Old” responses to foils. Participants completed the MST immediately before and after the intervention and again approximately 4 weeks after completing the intervention (average number of days post-intervention 31 ± 6). Thus, we evaluated changes pre- and post-intervention and also whether any observed changes in performance were still present 4 weeks after the completion of the exploration intervention.

**Figure 1.**
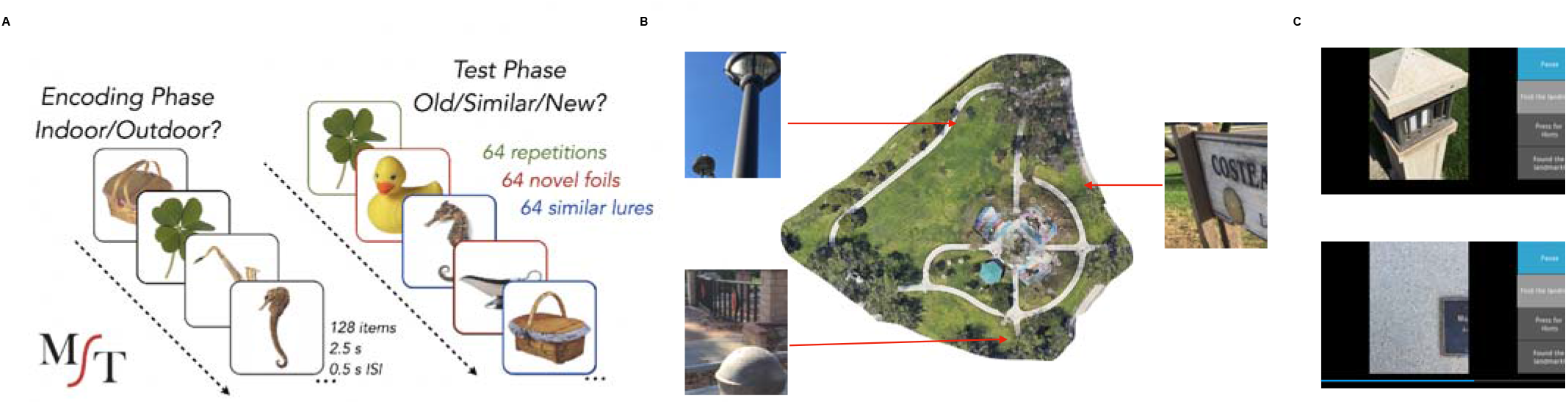
Behavioral task paradigms. A. Schematic of the Mnemonic Similarity Task. B. One of the parks used for the exploration task with three example landmarks and their locations within the park. C. Screenshots from the real-world exploration app.

#### 2.2.3 Real-World Exploration Task

Participants completed a scavenger hunt task in four small community parks (average of 3.3 ± 0.41 acres) in either Laguna Hills, CA or Irvine, CA. Participants were provided with a cell phone loaded with the scavenger hunt task application. The task required participants to complete scavenger hunts in four different parks, one park per week, 5 days per park (20 sessions total). There were 16 landmarks in each park (benches, statues, signs, etc.) and each day 8 of these landmarks were randomly selected (Figure 1B). Cues to landmarks were presented one at a time (zoomed in pictures of a component of the landmark so-as to hide any large contextual cues), and participants were instructed to find and walk to the landmark (Figure 1C). The app logged GPS coordinates once every second so that we could reconstruct the paths that were taken to each landmark. All participants were trained by the experimenter in how to use the phone and the app on the first day of the intervention. Participants completed the 20 sessions on their own schedule but were only allowed to do one session per day. The app included a hint function where, if participants were lost, they could tap a button and a bar appeared at the bottom of the screen which filled in from left to right indicating how close or far away they were from the landmark (Figure 1C bottom). They were told to use this if they had already adequately searched for and yet could not find the landmark for that trial.

## 3 Results

### 3.1 Excess Path

Statistics were performed using a combination of JASP, Prism 8 and Python 3. To estimate the amount and rate of learning that occurred over the time in the park, we calculated normalized excess path and heading error for each trial. Trials in which the GPS signal was lost, the app crashed, or there was clear user error (e.g., accidentally closing the app before trial completed) were removed (average 11 trials per participant). We then calculated the normalized excess path for each trial by (total path length – ideal path length)/ideal path length. The normalization allows us to compare the excess path of trials where the start of the trial was near the target to those where the target was far away from the start point. We averaged these excess path values across days and parks to control for variability in path due to the random selection of landmarks on each day. We entered average excess path over each of the 5 days into a repeated-measures one-way ANOVA and found a significant main effect of day (*F*_(4,96)_ = 11.60, *p* = .0000001) (Figure 2A). A planned contrast revealed a significant decrease in excess path from Day 1 to Day 5 (t_(24)_ = 4.896, *p* = .000027, *d* = .979). These results indicate that participants had learned the locations of the landmarks and that the routes taken to the landmarks became more efficient over time.

**Figure 2.**
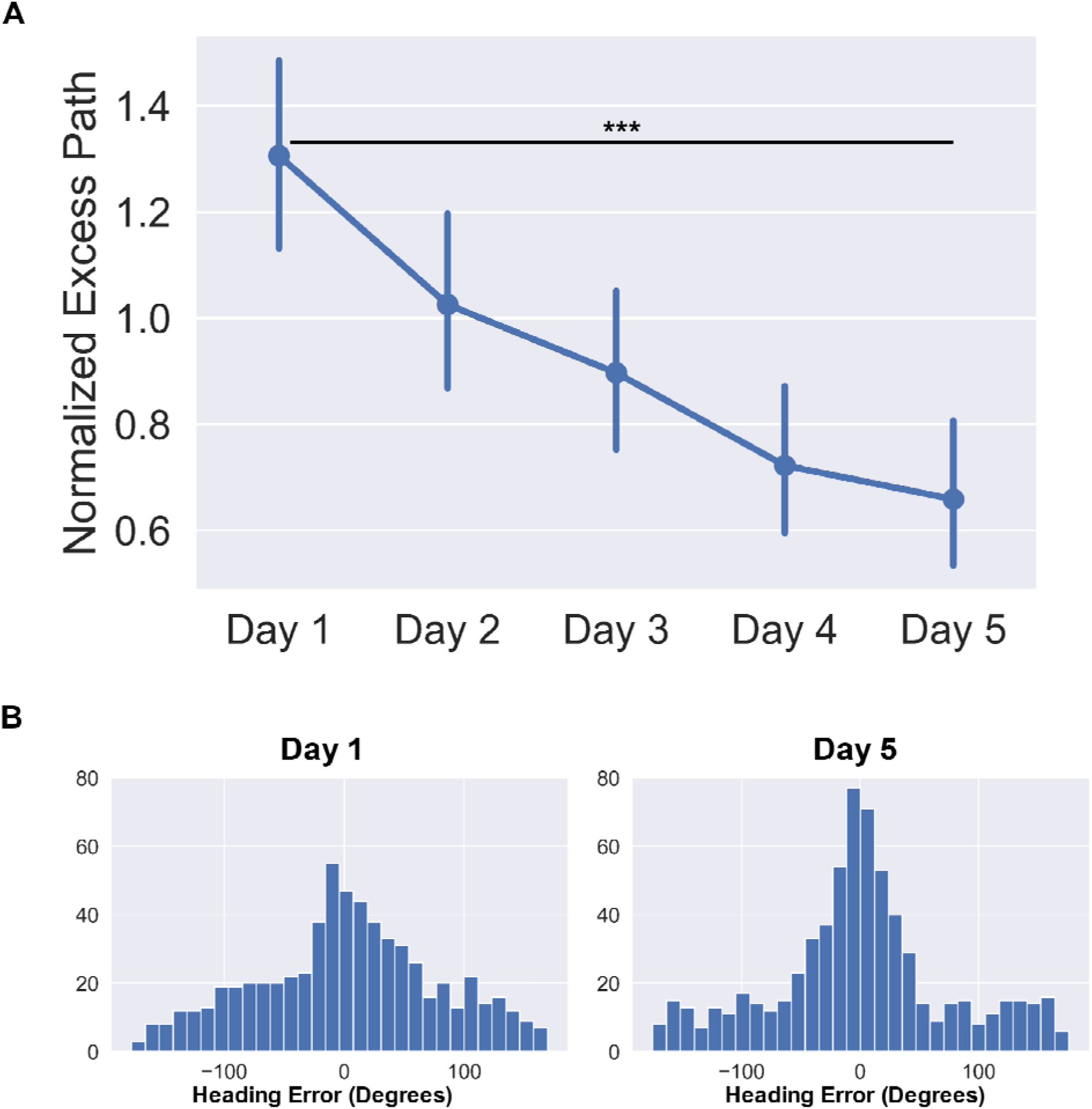
Behavioral data from the real-world exploration task. A. Average normalized excess path (±SEM) across all five days. B. Heading angle errors for day 1 and day five across the four parks. *** *p* < .001

### 3.2 Heading Error

To estimate of how well participants knew which direction to go immediately upon seeing the landmark for that trial, we computed errors in heading angle as the angular difference between the vectors from the start of the trial to the target and from the start of the trial to their position 10 seconds after the trial began. To determine if the distribution of errors were different from day 1 to day 5, we entered the distributions into a two-sample Kolmogorov-Smirnov test. We found that the distribution of heading error was significantly different on day 5 than it was on day 1 (*D* = .1075, *p* = .000171) (Figure 2B). Additionally, we modeled both day 1 and day 5 heading errors as mixture models of a von Mises distribution (circular version of a Gaussian distribution to model angular error in memory for the location) and a uniform distribution (to model random guesses). We computed the spread parameter (*k*) of the von Mises (akin to 1/σ of angular error) for both day 1 and day 5 heading errors which estimates how tightly the distribution is clustered around the mean. We compared the fits of two models, one which constrained the *k* parameters to be equal (*k*_day1_ = *k*_day5_) and one where the *k* parameters were allowed to differ between the two models (*k*_day1_ ≠ *k*_day5_). We found that the model in which the *k* parameters were allowed to differ significantly increased the quality of fit (*k*_day1_ = 1.98, *k*_day5_ = 5.95, difference in AIC = 10.45). These results again highlight the fact that participants learned the locations of the landmarks and by day 5 were heading in the right direction within the first 10 seconds of the trial.

### 3.3 Number of Hints

We logged the number of hints participants requested each day. For one participant, the number of hints per day was not accurately recorded, so they were removed from the analysis. We entered the number of hints requested across each of the five days in the park into a repeated-measures one-way ANOVA and found a significant main effect of day (*F*_(4,369)_ = 8.60, *p* = .0019). Importantly, post-hoc comparisons revealed a significant decrease in the number of hints from day 1 to day 5 (*t*_(24)_ = 3.619, *p* = .003, *d* = .619), again indicating that participants had learned the landmark locations and required fewer hints to find them by day 5. Overall, the results from the park show that participants had learned the locations of the landmarks in each park and the spatial relationship between them.

### 3.4 Mnemonic Similarity Task

Participants completed the MST both before and after the real-world exploration intervention, as well as at a follow-up appointment approximately four weeks after ending the intervention, with different sets of items used each time. We predicted that lure discrimination (LDI) would improve following the intervention, while recognition memory scores would not change owing to the differential degree of hippocampal involvement in the two measures. Four participants who performed significantly below chance on the MST’s recognition memory component were removed from the analysis. To test our hypotheses, we ran separate mixed model ANOVAs for recognition and LDI scores with Phase as a within-subjects factor (Pre-, Post, and Follow-Up) and Age as a between-subjects factor. For recognition, we observed no significant effect of Phase (*F*_(2,16)_ = .988, *p* = .394), no effect of Age (*F*_(12,8)_ = 1.309, *p* = .360), and no Age x Phase interaction (*F*_(24,16)_ = .666, *p* = .821) (Figure 3B). For LDI, we did find a significant main effect of Phase (*F*_(2,16)_ = 12.345, *p* = .00057) (Figure 3B). Bonferroni-corrected comparisons revealed a significant difference in LDI from pre-test to post-test (*t*_(20)_ = 5.238, *p_bonf_* = .00002, Cohen’s *d* = 1.143) as well as a significant difference pre-test to follow-up (*t*_(20)_ = 3.980 *p_bonf_* = .001, *d* = .868). There was no significant main effect of Age (*F*_(12,8)_ = 1.489, *p* = .292) or an Age x Phase interaction (*F*_(24,16)_ = 1.781, *p* = .118) (Figure 3C). These results indicate that, as we predicted, no change was observed in recognition memory, while LDI improved in our participants following the intervention and remained significantly higher than baseline approximately four weeks after finishing the intervention. The LDI improvement was also observed independent of age, indicating that, within the restricted range of this older sample, being younger does not confer a greater benefit from this training.

**Figure 3.**
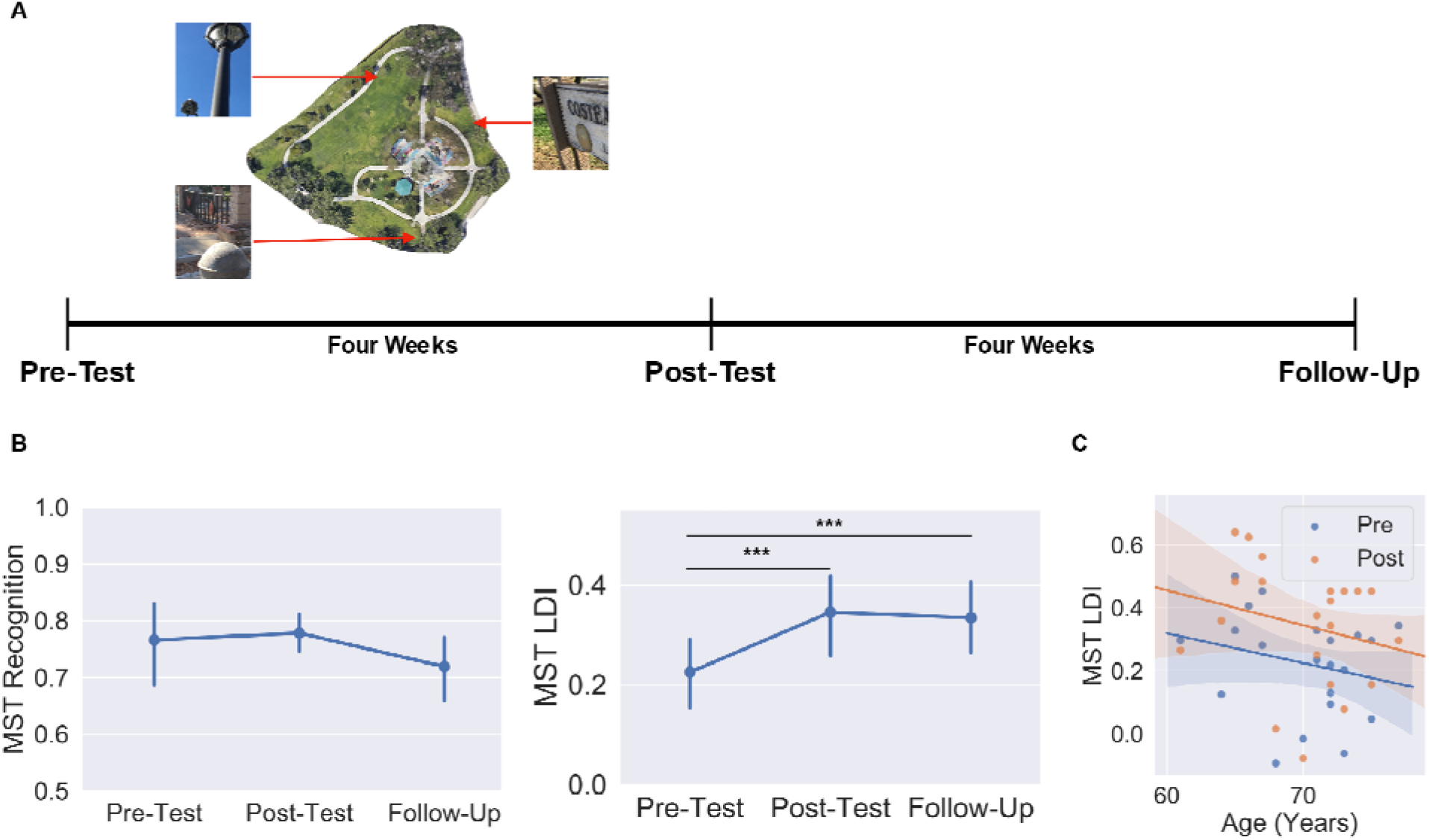
Schematic of behavioral testing timeline. A. Timeline used for behavioral testing. MST was administered at pre-test, post-test, and follow-up. B. Average recognition and LDI scores (±SEM) across the three testing sessions. C. Scatter plot of pre-test and post-test MST LDI scores by age. *** *p* < .001

### 3.5 MST and Park Performance

We next wanted to determine how performance in the parks was related to change in LDI. First, for each participant we computed the slope of their excess path over the five days in the park as a metric for how quickly they learned the landmark locations. We then correlated the slope with their change in LDI (post-pre). We found no significant relationship between excess path slope and LDI change (*r*_(20)_ = .210, *p* = .4517), indicating that improvement in lure discrimination was possible across many levels of exploration performance. We also failed to find a significant relationship between the average distance walked per day and change in LDI (*r*_(20)_ = −.116, *p* = .624), suggesting that our observed improvement in lure discrimination cannot be explained by the amount of physical activity. Finally, we wanted to test whether those who were performing better pre-intervention had the greatest benefit from the intervention by correlating preintervention LDI with LDI change (post-pre). We, again, found no relationship between change in LDI score and pre-intervention baseline (*r*_(20)_ = .1863, *p* = .3725). Collectively, these results highlight that benefits can be seen from our intervention regardless of performance level at baseline, how quickly one learns the target locations, or the amount of physical activity engaged in during the intervention. These are important aspects to consider when determining the efficacy of interventions for the larger population.

## 4 Discussion

We tested whether a real-world exploration intervention could improve hippocampal-based memory performance in healthy older adults. Participants completed 20 sessions of a scavenger hunt in local parks over 4 weeks. Memory performance was measured with the Mnemonic Similarity Task before and after the intervention, as well as at a four-week follow-up. Excess path measures decreased over time in the parks, as did initial heading error and the number of hints requested, indicating that participants were learning the locations of the landmarks in the park and were able to take more direct paths to them. Participants showed significant improvement in mnemonic discrimination (the MST’s pattern separation based LDI metric) but no significant change in recognition, supporting our hypotheses. Additionally, the boost in LDI was still seen at a four-week follow-up test. These results are in line with previous work showing improved pattern separation based memory in rodents (Creer, Romberg, Saksida, van Praag, & Bussey, 2010) as well as young adults (Clemenson et al., 2019; Clemenson & Stark, 2015) following environmental enrichment. The magnitude of LDI change in the current study (ΔLDI ≈ 0.1, d ≈ 0.8) is also in line with previous studies showing LDI change as a result of a behavioral intervention using video games as a proxy for environmental enrichment in both younger (Clemenson et al, 2019; Clemenson & Stark, 2015) and older adults (Clemenson, Stark, Rutledge, & Stark, under review). Additionally, these results support the role of spatial exploration as a viable component of enrichment for promoting changes in hippocampal dependent memory. Future studies should include a broader assay of cognition including tests of attention and executive function to assess performance changes in domains that are not necessarily hippocampally-dependent.

Investigating the role of environmental enrichment in humans is difficult since we already exist in an enriched environment. However, in rodents, one effective component of enrichment seems to be the active exploration of the environment (Freund et al., 2013), something we incorporated here and in our prior work in humans using video games (Clemenson et al., 2019; Clemenson & Stark, 2015). An important difference between the previous work using video games and the current study is that our participants were physically moving in and exploring the environment. Physical exercise has been linked to changes in hippocampal volume and improvements in memory performance in humans (Erickson et al., 2011), so it is important to address the role of exercise in a real-world exploration task compared to video game interventions. We saw similar improvements in memory ability as we previously found with a video game intervention, yet we saw no correlation between memory improvement and the average distance walked per day. We cannot conclude that the physical activity inherent in our intervention had no impact. We did, however, find that improvements in memory could be seen independent of the amount of exercise.

Another important finding from the current study is that the observed improvement in memory ability was not dependent on how well one learned the spatial locations during the exploration intervention. This observation is critical for designing interventions for the larger population, since there will be natural variability in how well participants can perform an exploration task. If the intervention only benefits those who are able to learn quickly or start at a higher level of baseline performance, then the intervention is not as useful for the general population. We should note, however, that it is almost certain that participants were incidentally encoding many aspects of the parks as part of their experience during the intervention. We do not know whether variance in how much was learned in these aspects would correlate with the observed improvements in memory ability. We also note that memory improvement was observed across all ages in our sample. The age range of our sample is limited, and it is possible that if including a larger age range, we may observe a relationship between participant age and the memory improvement. Given the current results however, it appears that our intervention can have equal benefit for older adults at any age.

Finally, though our intervention required a specific application to complete the scavenger hunt, our methods can be adapted to daily life quite easily. Often, older adults have highly routine schedules and habits that maybe require less spatial learning than would younger adults, who have more flexible and less routine day-to-day experiences. In so far as these results and our prior work support the general notion that enrichment improves hippocampal based memory performance, they suggest alternate avenues for inducing this enrichment. For example, learning the locations of the landmarks in the park and how to navigate between them could translate into taking new routes to the grocery store or a friend’s house. Learning new spatial relationships in this way could mimic the cognitive engagement required for this intervention and on a long-term basis could have significant benefits for hippocampal-dependent memory.

### Conclusion

We have shown that participating in a four-week real-world exploration intervention boosted performance on mnemonic pattern separation in older adults. These results show a remarkably similar pattern and magnitude of change on MST recognition and LDI scores as seen in our previous work that used video games to provide the spatial exploration and enrichment in younger participants (Clemenson et al., 2019; Clemenson & Stark, 2015) and in older participants (Clemenson et al., under review). Importantly, our intervention can be easily implemented into a daily schedule and potentially alleviate some of the age-related decline in hippocampal-dependent memory.

## Acknowledgements

This research was supported by a grant from the Dana Foundation. In addition, we would like to acknowledge additional salary support from NIA R21AG056145 and NIA R01AG034613. No conflicts of interest exist for any of the authors.

